# Maternal approach behaviors toward neonatal calls are impaired by mother’s experiences of raising pups with a risk gene variant for autism

**DOI:** 10.1101/2020.05.21.107540

**Authors:** Risa Kato, Akihiro Machida, Kensaku Nomoto, Gina Kang, Takeshi Hiramoto, Kenji Tanigaki, Kazutaka Mogi, Noboru Hiroi, Takefumi Kikusui

## Abstract

How the intrinsic sequence structure of neonatal mouse pup ultrasonic vocalization (USV) and maternal experiences determine maternal behaviors in mice is poorly understood. Our previous work showed that pups with a *Tbx1* heterozygous (HT) mutation, a genetic risk for autism spectrum disorder (ASD), emit altered call sequences that do not induce maternal approach behaviors in C57BL6/J mothers. Here, we tested how maternal approach behaviors induced by wild-type and HT USVs are influenced by the mother’s experience in raising pups of these two genotypes. The results showed that wild-type USVs were effective in inducing maternal approach behaviors when mothers raised wild-type but not HT pups. The USVs of HT pups were ineffective regardless of whether mothers raised HT or wild-type pups. However, the sequence structure of pup USVs had no effect on the general, non-directional incentive motivation of maternal behaviors. Our data show how the mother’s experience with a pup with a genetic risk for ASD alters the intrinsic incentive values of USV sequences in maternal approach behaviors.

## 1. Introduction

Autism spectrum disorder (ASD) is defined by deficits in social communication and interaction, as well as deficits in cognitive flexibility. While evidence supports a strong genetic influence on its etiology, the precise mechanisms through which genetic variation causally results in ASD are still poorly understood. Hemizygous deletion at human 22q11.2 is one of the rare copy number variants that are robustly associated with ASD (Niklasson et al., 2002; Fine et al., 2005; Antshel et al., 2007; Kates et al., 2007; Schneider et al., 2014; Hoeffding et al., 2017; Zinkstok et al., 2019). *TBX1* is one of the genes encoded in 22q11.2 and several cases of *TBX1* mutations are associated with ASD (Gong et al., 2001; Ogata et al., 2014; Paylor et al., 2006). The human TBX1 protein and its mouse ortholog (Tbx1) share a highly conserved amino acid sequence. In mice, *Tbx1* heterozygosity induces many behavioral phenotypes relevant to ASD, including defects in social interaction, social communication, and working memory (Hiramoto et al., 2011; Takahashi et al., 2016; Hiroi and Yamauchi, 2019).

In humans, cries in incipient ASD babies are atypical and such cries are not well understood or are more negatively perceived by mothers (Esposito et al., 2017; Esposito & Venuti, 2008, 2010a, 2010b). We previously showed that *Tbx1* heterozygous (HT) pups emit atypical sequences of ultrasonic vocalizations (USVs) (Hiramoto et al., 2011). Using a procedure that evaluates maternal approach behaviors evoked by played-back pup USVs (Mogi et al., 2017; Okabe et al., 2010, 2013; Okabe, Nagasawa, Mogi, & Kikusui, 2017), we showed that USVs of *Tbx1* HT pups did not elicit maternal approach behaviors in C57BL/6J mothers (Takahashi et al., 2016), suggesting that HT pup USVs lack incentive values to elicit maternal behavior.

Given that the sequence structure of pup USVs is a determinant of maternal behaviors, we designed an experiment to examine how the experience of a mother with her own WT and HT pups subsequently determines her response to the USVs of other WT and HT pups. To control for USV familiarity and strictly test the sequence structure of USVs, we used WT and HT pup USVs of different mothers previously collected (Hiramoto et al., 2011). Our data show that the incentive value of USV is differentially modified by prior experience of mothers with pups of different genotypes.

## 2. Materials and Methods

### 2-1. Mouse

Congenic *Tbx1* HT and WT mice were generated from breeder pairs of congenic *Tbx1* HT male mice (Hiramoto et al., 2011) and WT female mice. Our mice were backcrossed to C57BL/6JJcl mice (CLEA Japan, Inc., Japan) for more than 10 generations to control for genetic background (Hiroi, 2018). The mice were maintained at a constant temperature (24 ± 1 °C) and humidity (45 ± 5 %) under a 12 h-12 h light-dark cycle (lights off from 18:00 to 06:00). They were housed in standard cages (175 × 245 × 125 mm) containing approximately 3 mm grain-size corn cob bedding (“Shepherd’s Cob,” Shepherd Specialty Papers, USA) and nesting material (“Parumasu ¼,” Material Research Center Co., Ltd., Japan) with *ad libitum* access to food and water.

### 2-2. Equipment for playing back recorded pup USVs

We used *Tbx1* HT and WT pup USVs recorded in our previous study (Takahashi et al., 2016). To control for familiarity of USVs and strictly test the sequence structure of WT and HT USVs, we did not use the USVs of pups that were raised by their mothers. We used an nc-Si emitter (Tokyo University of Agriculture and Technology) composed of a surface-heating thin-film electrode, nanocrystalline silicon (ns-Si) layer, and single-crystalline silicon wafer. We used RECORDER NI-DAQMX to play back the recorded sounds. A power amplifier (Asbir, Japan) was connected to a vocalization-analyzer system (MK-1500, Muromachi Kikai Co., Ltd., Japan) (Mogi et al., 2017; Okabe et al., 2010, 2013; Okabe, Nagasawa, Mogi, & Kikusui, 2017). The peak frequency and peak amplitude were not different between WT and HT pup USVs (Takahashi et al., 2016).

### 2-3. Test cage

The apparatus and procedure for playing back pup USVs and evaluating maternal approach behaviors were identical to those described in our previous study (Takahashi et al., 2016). The test apparatus was a standard Plexiglas mouse home cage (175 × 245 × 125 mm) with two tubes (diameter of 40 mm, length of 150 mm; see **Suppl. Figure 1a**). Both long sides of the cage had 12 slits (8 × 12 mm, 4 mm between slits, 2 cm from the right corner, and 1 cm from the bottom) to reduce echoing of sound inside the apparatus. The test apparatus contained novel bedding and was placed in a soundproof box (Muromachi Kikai Co., Ltd., Japan, 705 × 795 × 725 mm; the inner wall was covered with sponge) in a soundproof room (Kawai Acoustic System Co., Ltd., Japan) under a red lamp. The odor of the mother’s own pups, in the form of bedding and cotton smeared with bedding, was placed between the emitter and the mesh at the end of both tubes. One nc-Si emitter was placed facing the mesh-covered end of one of the two tubes (sound tube) and another without any cable connection was placed at the end of the other tube (no sound tube) (see **Suppl. Fig. 1**).

### 2-4. Procedure and analysis of maternal approach behaviors toward pup USV

In the present experiment, only WT mothers were examined. All of them were first-time mothers. Virgin WT female breeder mice, at 8-13 weeks old, were paired with HT male breeder mice for up to two weeks, during which male breeders were removed as soon as female breeders were identified as pregnant. The genotype of all pups was determined by polymerase chain reaction on the day when they were born and only four pups of the same genotype were kept with their own mother until they were weaned at PD 28. At 5-7 days after giving birth to pups, lactating WT mothers were used for behavioral testing with USVs of other pups. Each of eleven WT mothers raised four WT pups; each of the other WT mothers (n=9) raised four HT pups. The ratio of male to female pups randomly varied from litter to litter.

We used the previous USV recordings of 7-8 day old female pups that were closest to the average of each genotype group (Takahashi et al., 2016); these calls were not those of pups whose mothers were tested in the present study. We did not match WT and HT calls in terms of their acoustic parameters, because the group averages of WT and HT pup calls significantly differed in terms of frequencies, durations and sequence structures (Takahashi et al. 2016).

There were two experimental groups: mothers that raised WT pups and exposed to WT and HT pup USVs, and mothers that raised HT pups and exposed to WT and HT pup USVs. On Day 1, the home cage of a WT mother, with her own pups, was placed for 24 hours in the soundproof room where the experimental apparatus was stationed. Days 2 and 3 were test days. On Day 2, after a mother was placed in the test apparatus in the soundproof box for 30 minutes under the red lamp (habituation), she was tested with pup USVs for 5 minutes (test) (**Suppl. Figure 1b**). On Day 3, the mother was similarly tested with the pup USVs that were not used on Day 2. The order of pup USV presentation was counterbalanced: half of the mothers were presented with WT pup USVs on Day 2 and *Tbx1* HT pup USVs on Day 3; the opposite order of presentation was used for the other half (**Suppl. Figure 1c**). During the test session, the mother was allowed to freely move in the test apparatus. After each testing, she was returned to the home cage that was placed in the soundproof room.

The mother’s behavior was recorded by a video camera and analyzed for the time spent peeking (the mother’s upper body entered the entrance of each tube), time spent in the tube (the mother’s body including hind paws entered each tube), and time spent at the mesh (the mother touched the mesh at the end of each tube). Data were analyzed with time spent in the sound and no-sound tubes for each behavior.

### 2-5. Statistical analyses

As the homogeneity of variance and normality were violated in some cases, the entire data set was analyzed using non-parametric Wilcoxon tests. All statistical analyses were executed with SciPy in Python 3.6.9. *P* value < 0.05 was considered significant. One data set was excluded from analyses, because one mother escaped from the test cage during the Day 2 test session when WT pup USVs were played back.

## 3. Results

We found that, compared to the no-sound tube, WT mothers that raised WT pups spent more time in peeking at and staying in the sound tube from which WT pup USVs were played back (**Figure 1ab, Raised WT pups, WT USVs**); they did not stay longer at the mesh of the sound tube, compared to the no-sound tube (**Figure 1c, Raised WT pups, WT USVs**). Those mothers did not show any preference for HT pup USVs in any of the three measures (**Figure 1abc, Raised WT pups, HT USVs)**. Mothers that raised HT pups did not prefer WT or HT pup USVs in any of the three measures (**Figure 1abc, Raised HT pups**). We consider exploratory behaviors directed toward the source of an incentive stimulus as reflecting directional incentive motivation.

**Figure 1.**
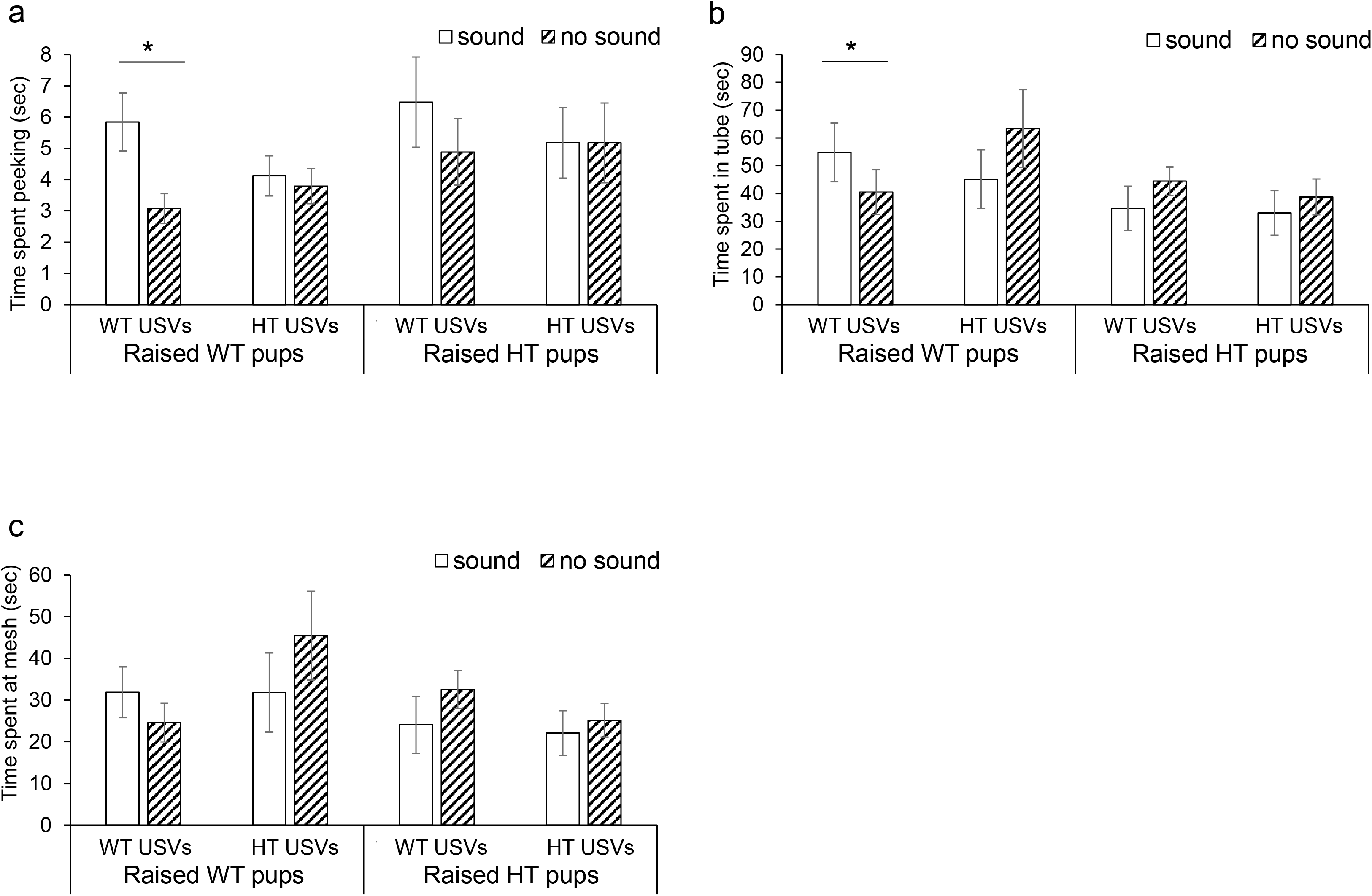
The average amount of time (+SEM) mothers spent in the sound and no-sound tubes is shown. When WT pup USVs were played back, WT mothers that raised WT pups preferred the sound tube to the no-sound tube in peeking at the tube entrance (**Raised WT pups WT USVs**, **a**, T=7.0, p = 0.037) and staying in the tube (**b**, T=7.0, p = 0.037); they did not show statistically significant preference in staying at the mesh (**c**, T=18.0, p = 0.33). When HT pup USVs were played back, WT mothers that raised WT pups did not show preference to the sound tube (**Raised WT pups HT USVs**, **a**, T=32.0**, p =**0.929; **b**, T=23.0, p =0.374; **c,** T=23.0, p =0.374). When WT pup USVs were played back, WT mothers that raised HT pups showed no preference (**Raised HT pups WT USV**, **a**, T=13.0, p =0.260; **b**, T=15.0, p =0.374; **c**, T=14.0, p =0.314). When HT pup USVs were played back, WT mothers that raised HT pups showed no preference (**Raised HT pups HT USVs**: **a**, T=22.0, p =0.953; **b**, T=15.0, p = 0.374; **c**, T=21.0, p = 0.859). \**p*<0.05, significant difference at 5% level, as determined by Wilcoxon non-parametric tests. Sound, a sound tube from which pup USVs were played back; no-sound, the tube from which no pup USVs were played back. Raised WT, WT USV; N=10; Raised WT, HT USV; N=11; Raised HT, WT USV; N=9; Raised HT, HT USV; N=9.

The total amount of time spent in both sound and no-sound tubes is an index of how motivated mothers were in searching for pups, independent of their accuracy in choosing the sound tube. When a mother’s search behavior is invigorated, but such exploratory behaviors are not directed toward the sound source, we define it as reflecting elevated levels of non-directional incentive motivation. This aspect of incentive motivation can be indexed by the total amount of time spent in both sound and no-sound tubes. Mothers spent indistinguishable amounts of total time in peeking at the tube entrances, staying in the tubes and at the mesh ends, regardless of whether they raised WT or HT pups or were tested with WT or HT pup USVs (**Suppl. Figure 2**).

## 4. Discussion

In the present study, we tested how maternal approach behaviors induced by WT and HT pup USVs are influenced by mother’s experience in raising pups of these two genotypes. The subject WT mothers were divided into two groups: one group of mothers raised WT pups and the other group of mothers raised HT pups. Both groups were sequentially tested with WT and HT pup calls in a counterbalanced order. Our data showed that 1) HT pup USVs lacked the incentive value for inducing directional maternal approach behaviors regardless of mothers’ experiences in raising HT or WT pups; 2) the experience of raising HT pups eliminated the directional incentive value of WT pup USVs to evoke maternal approach behaviors; 3) mothers that raised HT pups did not show directional maternal behaviors in response to HT pup USVs; and 4) mothers were equally motivated to non-directionally search for pups in response to USVs of any genotype regardless of the genotype of pups mothers raised. These observations could be interpreted as indicating that the directional incentive values and generalized, non-directional incentive values of USVs are dissociable and the former, but not the latter, is impacted by the sequence structure of USVs and the mother’s prior experiences.

A noteworthy technical aspect of the current study is that we did not use USVs of WT and HT pup of the mothers tested in this experiment. Instead, we used the USVs that were statistically closest to the genotype average of WT and HT pups of C57BL/6J background recorded in the previous study (Takahashi et al., 2016). This experimental design was needed to isolate the common pup USV sequences and control for simple familiarity of individually variable components of pup USV sequences. That so-selected WT USVs induce directional maternal approach in WT mothers suggests the existence of a common sequence structure among WT pups with a C57BL/6J strain genetic background, which is sufficient for directional maternal approach behavior even if the mother has never been exposed to such individual WT pups and their calls; HT pup USVs lacked such properties. Complementing this, we previously showed that C57BL/6J females that had their own C57BL/6J pups demonstrated maternal approach behaviors toward WT pup USVs but not toward *Tbx1* HT pup USVs, despite the fact that C57BL/6J mothers that were used for testing were never exposed to those USVs and never raised *Tbx1* HT or WT pups whose neonatal USVs were used for testing (Takahashi et al., 2016). Similarly, in Institute of Cancer Research (ICR) mice, mothers innately approach pup USVs of the same mouse strain (Okabe et al., 2010) and their own pups’ USVs (Mogi et al., 2017).

HT USVs did not induce directional maternal approach behaviors in mothers that raised WT pups. It could be due to the fact that WT mothers did not interact with HT pups and thus were not exposed to HT USVs in their home cages. This is unlikely, as experience of raising HT pups did not enable mothers to respond to HT pup USVs. However, WT mothers may simply require longer or more experiences with HT pups for HT USVs to evoke maternal behaviors. In any case, the relative directional incentive values of WT and HT pup USVs clearly differ.

A novel observation of the current study is that when mothers raised HT pups, WT USVs became ineffective in inducing directional maternal approach behaviors. This observation could be interpreted as indicating that HT pup or their USVs in home cages did not foster the mother’s ability to locate and prefer WT pup USV. WT mothers may have an innate neural circuit to respond to the common sequence elements of WT pup USVs, which are absent in HT pup USVs.

Some brain regions might be functionally required for processing the incentive value of pup USVs. Regions that convey information of pup USVs in the rodent mother’s brain might include various cortical and limbic structures, including the auditory cortex, basolateral amygdala, medial preoptic area of the hypothalamus, bed nucleus of the stria terminalis, and central nucleus of the amygdala (Okabe et al. 2013). In the auditory cortex of mothers but not naïve virgins, pup USV detection is improved by increasing the contrast between neural activity evoked by pup USVs and that evoked by background noise (Lin et al., 2013). The auditory cortex exhibit plastic changes by the interaction with pups; spike timing precision of the auditory cortex increases after experience with pups during heightened cortical oxytocin levels (Marlin, Mitre, D’Amour, Chao, & Froemke, 2015). Such plasticity might underlie different maternal behaviors in response to typical and atypical USVs. The deficits in directional maternal approach behavior toward WT pup USVs in mothers that raised *Tbx1* HT pups might be due to defective plastic changes in the brain by atypical USVs during mother-pup interactions. More work is needed to critically evaluate neuronal substrates through which the sequence structure of USVs contributes to maternal behaviors.

In conclusion, the present study showed how maternal experiences impact subsequent maternal incentive motivation in response to USVs. As experience with pups possessing an ASD risk gene variant negatively modifies maternal approach behavior toward USVs in mice, our data lend further support for the idea that the phenotypes of ASD risk gene carriers act as a new negative environmental factor for mothers, and further exacerbate, as a modifier, the consequence of genetic risk factors in the trajectory of ASD (Kikusui & Hiroi, 2017). These mouse observations might have translational values, as human mothers negatively perceive atypical vocalization of even their own incipient ASD babies (Esposito et al., 2017).

## Acknowledgments

The research reported in this publication was partly supported by the National Institutes of Health (R01MH099660 and R01DC015776 to NH) and JSPS KAKENHI (19K22373, 18H04890 to T.K. and 19K21822 to K.M.)

---

**Supplemental Figure 1**. The apparatus (a), procedure (b) and group assignment (c) of maternal approach behaviors. The order of WT and HT USV presentations was counter-balanced. Half mothers received WT USVs on Day 2 and HT USVs on Day 3; the other half received HT USVs on Day 2 and WT USVs on Day 3.

**Supplemental Figure 2.** The averages of the total time (+SEM) mothers spent in both sound and no-sound tubes are shown when WT USVs or HT USVs were played back. The total amount of time spent in both the sound and no-sound tubes is an index of how motivated mothers were in searching for pups, independent of their accuracy in choosing the sound tube. a. Time spent in peeking at the tube entrance, **Raised WT pups**, T=20.0, p = 0.445; **Raised HT**, T=20.0, p = 0.767; b. Time spent staying in the tube, **Raised WT pups**, T=20.0, p = 0.476; **Raised HT**, T=19.0, p = 0.678; c. Time spent staying at the mesh, **Raised WT pups**, T=15.0, p = 0.203; **Raised HT**, T=16.0, p = 0.441). As the assumption of normality was violated, we used non-parametric tests. Because each subject yielded the total time in response to WT and HT USVs, we used Wilcoxon non-parametric tests. Raised WT, WT USV; N=10; Raised WT, HT USV; N=11; Raised HT, WT USV; N=9; Raised HT, HT USV; N=9.

