## Supplementary figures and images for "Maternal approach behaviors toward neonatal calls are impaired by mother’s experiences of raising pups with a risk gene variant for autism"

### Supplemental Materials

Suppl. Figure 1

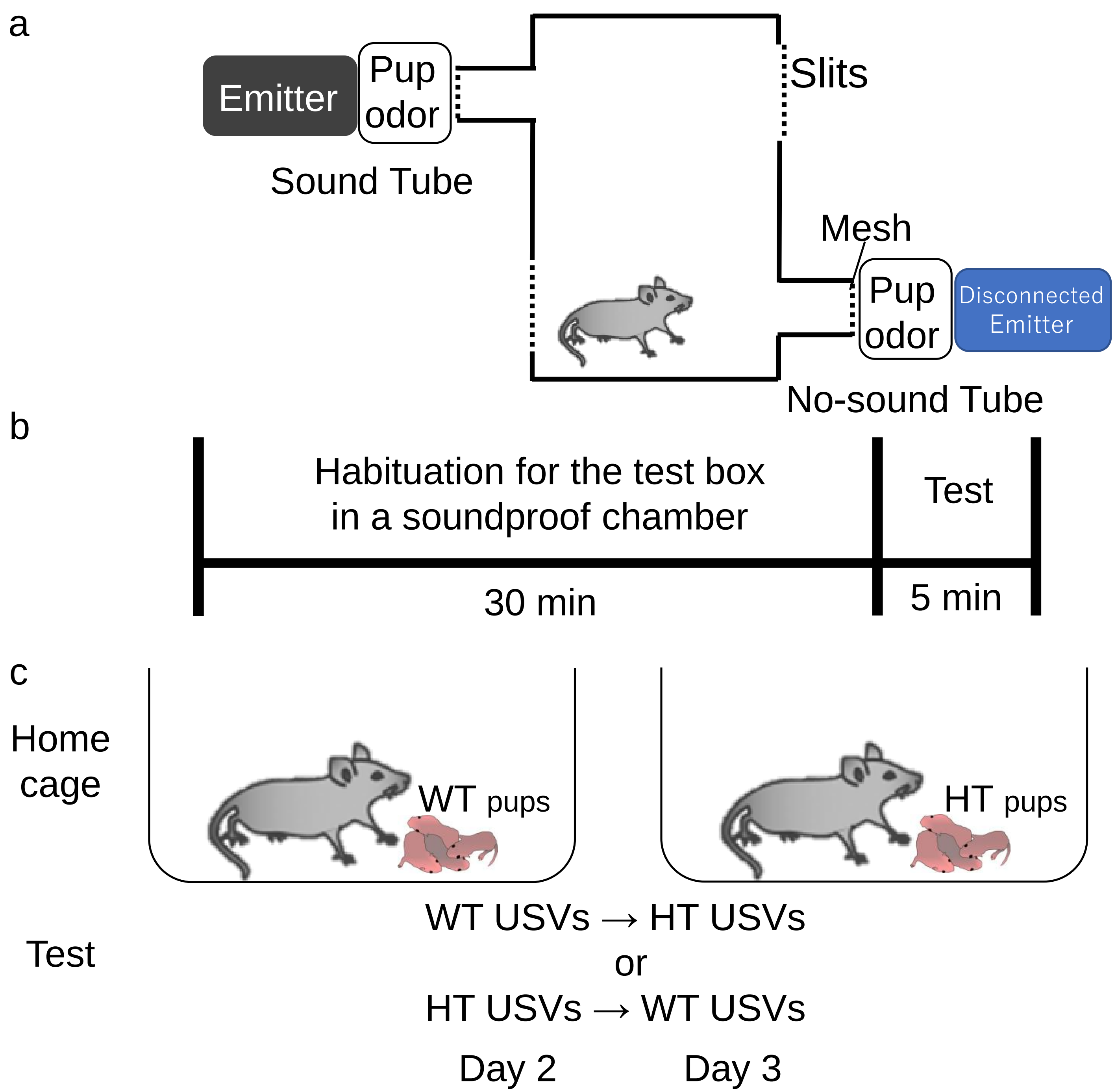

Suppl. Figure 2

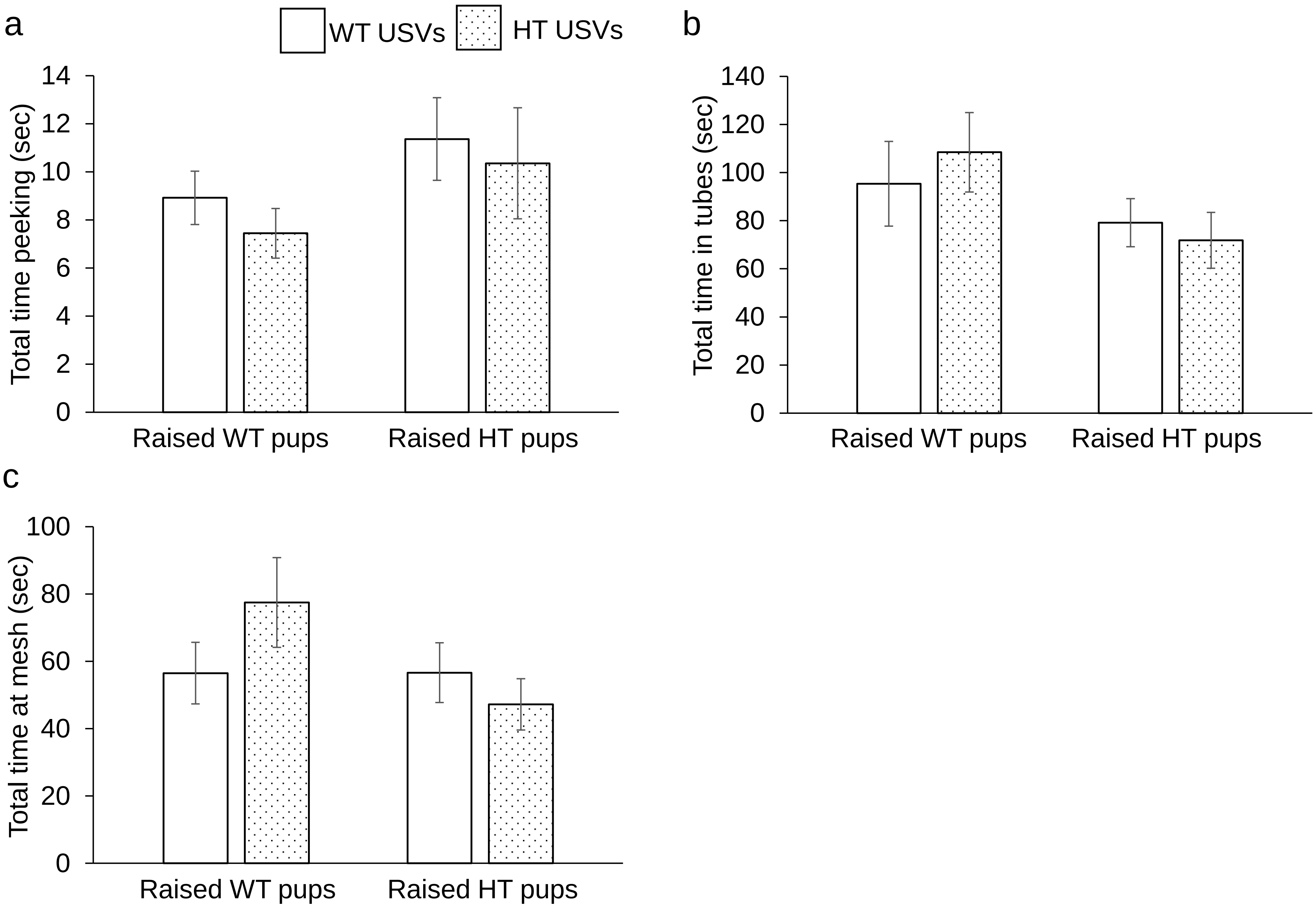
